# Novel Evolution of the Mineralocorticoid Receptor in Humans compared to Chimpanzees, Gorillas and Orangutans

**DOI:** 10.1101/2023.12.08.570861

**Authors:** Yoshinao Katsu, Jiawen Zhang, Michael E. Baker

## Abstract

We identified five distinct full-length human mineralocorticoid receptor (MR) genes containing either 984 amino acids (MR-984) or 988 amino acids (MR-988), which can be distinguished by the presence or absence of Lys, Cys, Ser, Trp (KCSW) in their DNA-binding domain (DBD) and mutations at codons 180 and 241 in their amino terminal domain (NTD). Two human MR-KCSW genes contain either (Val-180, Val-241) or (Ile-180, Val-241) in their NTD and three human MR-984 genes contain either (Ile-180, Ala-241) or (Val-180, Val-241) or (Ile-180, Val-241). Human MR-KCSW with (Ile-180, Ala-241) has not been cloned. In contrast, chimpanzees contain four MRs: two MR-988s with KCSW in their DBD or two MR-984s without KCSW in their DBD. Chimpanzee MRs only contain (Ile180, Val-241) in their NTD. A chimpanzee MR with either (Val-180, Val-241) or (Ile-180, Ala-241) in the NTD has not been cloned. Gorillas and orangutans each contain one MR-988 with KCSW in the DBD and one MR-984 without KCSW, and these MRs only contain (Ile-180, Val-241) in their NTD. A gorilla MR or orangutan MR with either (Val-180, Val-241) or (Ile-180, Ala-241) in the NTD has not been cloned. Together, these data suggest that human MRs with (Val-180, Val-241) or (Ile-180, Ala-241) in the NTD evolved after humans and chimpanzees diverged from their common ancestor. Considering the multiple functions in human development of the MR in kidney, brain, heart, skin, and lungs, as well as MR activity in interaction with the glucocorticoid receptor, we suggest that the evolution of human MRs that are absent in chimpanzees may have been important in the evolution of humans from chimpanzees. Investigation of the physiological responses to corticosteroids mediated by the MR in humans, chimpanzees, gorillas and orangutans may provide insights into the evolution of humans and their closest relatives.

## 1. Introduction

The mineralocorticoid receptor (MR) is a ligand-activated transcription factor, belonging to the nuclear receptor family, a diverse group of transcription factors that arose in multicellular animals [1–5]. The traditional physiological function of the MR is to maintain electrolyte balance by regulating sodium and potassium transport in epithelial cells in the kidney and colon [6–10]. In addition, the MR has important physiological functions in many other tissues, including brain, heart, skin and lungs [10–17].

The MR and its paralog, the glucocorticoid receptor (GR), descended from an ancestral corticoid receptor (CR) in a cyclostome (jawless fish) that evolved about 550 million years ago at the base of the vertebrate line [18–25]. A descendent of this ancestral steroid receptor, the CR in lamprey (*Petromyzon marinus*), is activated by aldosterone [26,27] and other corticosteroids [26,27]. Lampreys contain two CR isoforms, which differ only in the presence of a four amino acid insert Thr, Arg, Gln, Gly (TRQG) in their DNA-binding domain (DBD) [28] (Figure 1). We found that several corticosteroids had a similar half-maximal response (EC50) for lamprey CR1 and CR2 [27]. However, these corticosteroids had a lower fold-activation of transcription for CR1, which contains the four amino acid insert, than for CR2 suggesting that the deletion of the four amino acid sequence in CR2 selected for increased transcriptional activation by corticosteroids of CR2 [27,28].

**Figure 1.**
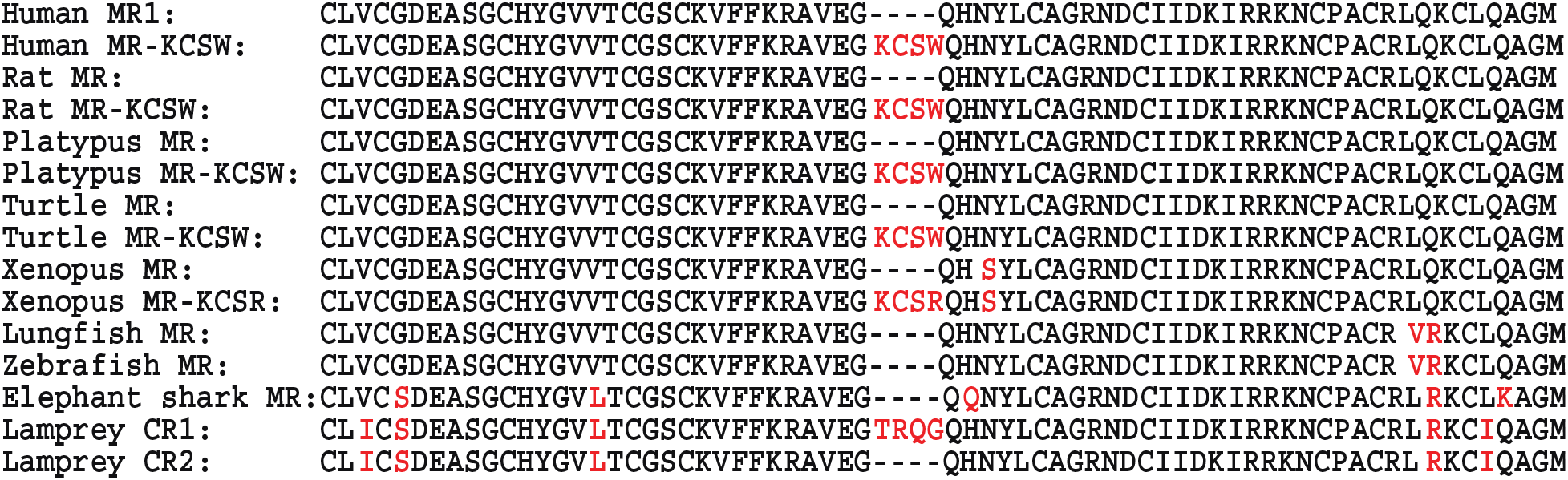
Comparison of the DNA-binding domain on human MR1, human MR-KCSW, rat MR, rat MR-KCSW, Platypus MR, Platypus MR-KCSW, Turtle MR, Turtle MR-KCSW, *Xenopus* MR, *Xenopus* MR-KCSR, lungfish MR, zebrafish MR, elephant shark MR, and lamprey CR1 and CR2.

A distinct MR and GR first appear in sharks and other cartilaginous fishes (Chondrichthyes) [20,22,29–32]. The DBD in elephant shark MR and GR lacks the four amino acid sequence found in lamprey CR1 [27] (Figure 1). We inserted this four-residue sequence from lamprey CR1 into the DBD in elephant shark MR and GR and found that in HEK293 cells co-transfected with the TAT3 promoter, the mutant elephant shark MR and GR had lower transcriptional activation by corticosteroids than did their wild-type elephant shark MR and GR counterparts, indicating that the insertion of the four amino acid sequence into the DBD of wild-type elephant shark MR and GR had a similar effect on transcriptional activation as the KCSW insert had in the DBD of lamprey CR1 [28].

Based on these results with lamprey CR1 and CR2, we analyzed the DBD sequence of human MR, which had been cloned, sequenced and characterized by Arriza et al. [33], and we found that like elephant shark MR, this human MR (MR1) lacks a four-residue segment in its DBD. This human MR has been widely studied [8,10,12,13,34–36]. Unexpectedly, our BLAST [37] search with the DBD from this human MR found a second, previously described, human MR with a KCSW insert (MR-KCSW) in its DBD [38–40] (Figure 1). A homology model of human MR with KCSW in the DBD constructed by Wickert et al. [40] found no distortion by the four amino acids on adjacent secondary structures of the DBD in human MR, consistent with activation of this MR by corticosteroids.

As described later, further BLAST searches found two full-length human MRs with this insert (MR-KCSW) and three full-length human MRs without this insert. The three human MRs without the KCSW insert contain either (Ile-180, Ala-241) or (Val-180, Val-241) or (Ile-180, Val-241) in their amino terminal domain (NTD) (Figure 2). The two human MR-KCSW splice variants contain either (Val-180, Val-241) or (Ile-180, Val-241) in their NTD (Figure 2). A human MR-KCSW with (Ile-180, Ala-241) has not been cloned.

**Figure 2.**
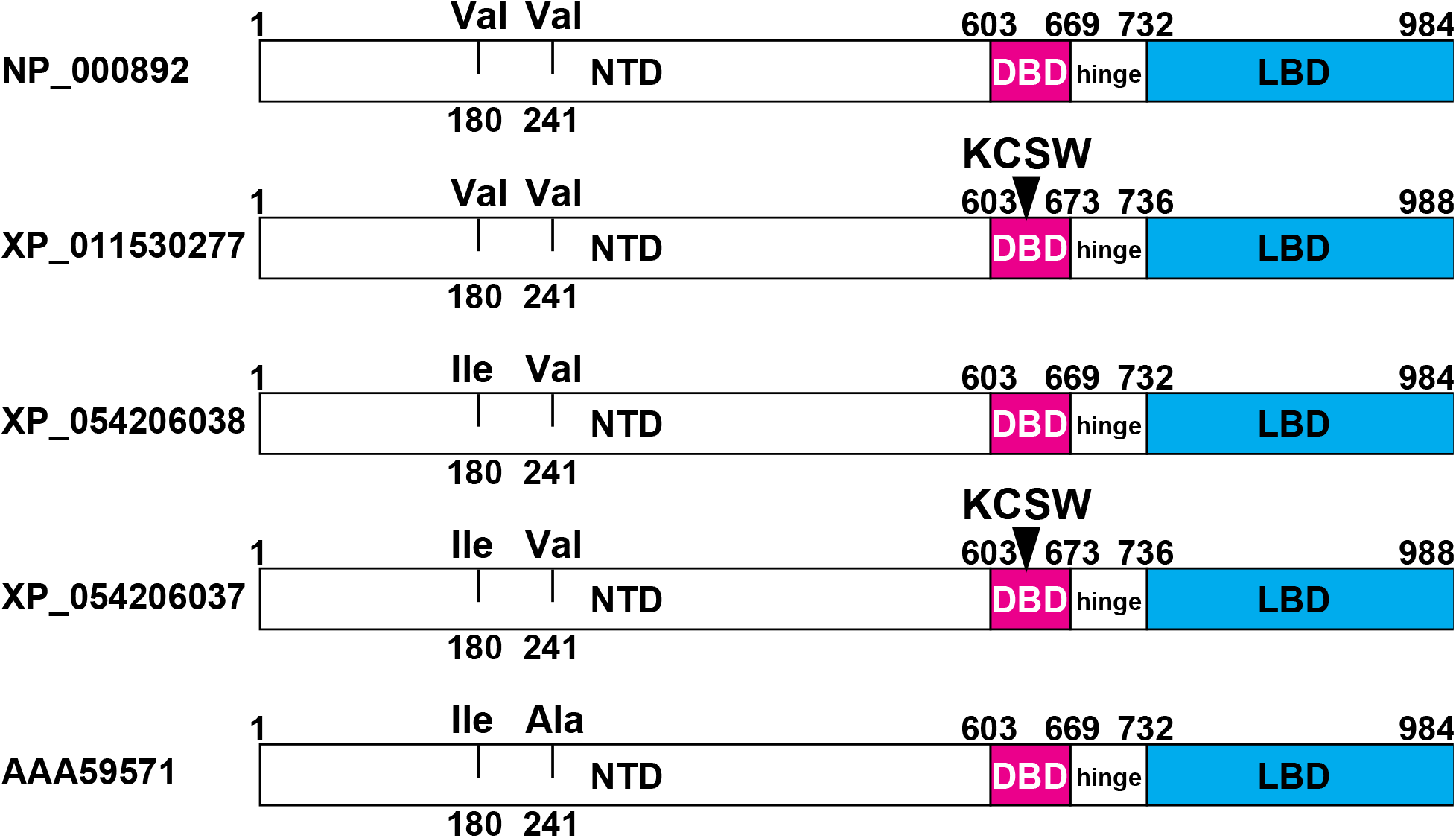
Comparison of five full-length human mineralocorticoid receptors. There are three human MRs with 984 amino acids. They contain either valine-180 and valine-241, isoleucine-180 and valine-241 or isoleucine-180 and alanine-241 in the amino-terminal domain (NTD). There are two human MRs with 988 amino acids and KCSW in the DBD and either valine-180 and valine-241 or isoleucine-180 and valine-241 in the NTD. A human MR with KCSW and isoleucine-180 and alanine-241 in the NTD was not found in GenBank.

Here, in this brief report, we describe our evolutionary analysis of human MR from a comparison of five full-length human MRs with four full-length MRs in chimpanzees and two full-length MRs in gorillas, and orangutans. We find that chimpanzees, gorillas, and orangutans lack an MR with either (Ile-180, Ala-241) or (Val-180, Val-241) in their NTD. We propose that MRs with these amino acids in the NTD evolved in humans after divergence of humans from chimpanzees. Due to the multiple functions in human development of the MR alone [11,15,16,41], as well as due to the interaction of the MR with the GR [17,42–46], we suggest that the evolution of three distinct MRs in humans that are absent in chimpanzees may have been important in the evolution of humans from chimpanzees.

The DBD of the human MR-KCSW splice variant has an insertion of four amino acids that is absent in human MR1. Otherwise, the rest of the sequences of human MR and human MR-KCSW are identical. Differences between the DBD sequence in human MR and selected vertebrate MRs are shown in red. Protein accession numbers are AAA59571 for human MR, XP_011530277 for human MR-KCSW, NP_037263 for rat MR, XP_038953451 for rat MR-KCSW, XP_007669969 for platypus MR, XP_016083764 for platypus MR-KCSW, XP_043401725 for turtle MR, XP_037753298 for turtle MR-KCSW, NP_001084074 for *Xenopus* MR, XP_018098693 for *Xenopus* MR-KCSR, BCV19931 for lungfish MR, NP_001093873 for zebrafish MR, XP_007902220 for elephant shark MR, XP_032811370 for lamprey CR1, and XP_032811371 for lamprey CR2.

## 2. Methods

The Basic Alignment Search Tool (BLAST) [37,47] was used to search GenBank with the sequence of human MR (Accession AAA59571) for similar proteins in humans, chimpanzees, gorillas and orangutans. We also retrieved MR sequences from rat and platypus, as well as two basal vertebrates: *Xenopus laevis* and turtles. We also retrieved MR sequences from ancestors of terrestrial vertebrates: lungfish, elephant shark and lamprey. The DBDs of these vertebrates are shown in Figure 1.

## 3. Results and Discussion

### 3.1 Novel Mineralocorticoid Receptors in Humans

A BLAST [37,47] search of GenBank retrieved five distinct full-length human MR genes (Figure 2), which are identical in the LBD and hinge segment, but differ in the DBD and NTD. Two full-length human MRs, with 988 amino acids, contain a KCSW insert in the DBD (Figure 2), which is absent in the human MR with 984 amino acids, cloned by Arriza et al. [33] (Figure 2), as well as two other full-length MRs with 984 amino acids (Figure 2). These three human MRs with 984 amino acids contain either (Ile-180, Ala-241), (Val-180, Val-241) or (Ile-180, Val-241) in the NTD (Figure 2). The two human MR-KCSWs contain either (Val-180, Val-241) or (Ile-180, Val-241) in their NTD. A human MR-KCSW sequence with (Ile-180, Ala-241) has not been deposited in GenBank.

### 3.2 Chimpanzees Have Four Full-Length Mineralocorticoid Receptors

Our BLAST search retrieved four full-length chimpanzee MRs (Figure 3). Two of these chimpanzee MRs have DBDs that are identical to the DBD in human MRs without the KCSW insert (Figure 1, Figure 3) and two chimpanzee MRs have DBDs that are identical to the DBD in human MR-KCSW (Figure 1, Figure 3). All four chimpanzee MRs contain (Ile-180, Val-241) in their NTD, which is similar to the NTD in one human MR (Figure 2). Chimpanzees lack an MR with either (Ile-180, Ala-241) or (Val-180, Val-241) in their NTD. The different chimpanzee MR sequences contain either a Ser-591 or an Asn-591 in their NTD (Figure 3). Human MRs contain a Ser-591 correspondingSer-591 in chimpanzee MR. Our BLAST search did not find a human MR with Asn-591 in the NTD.

**Figure 3.**
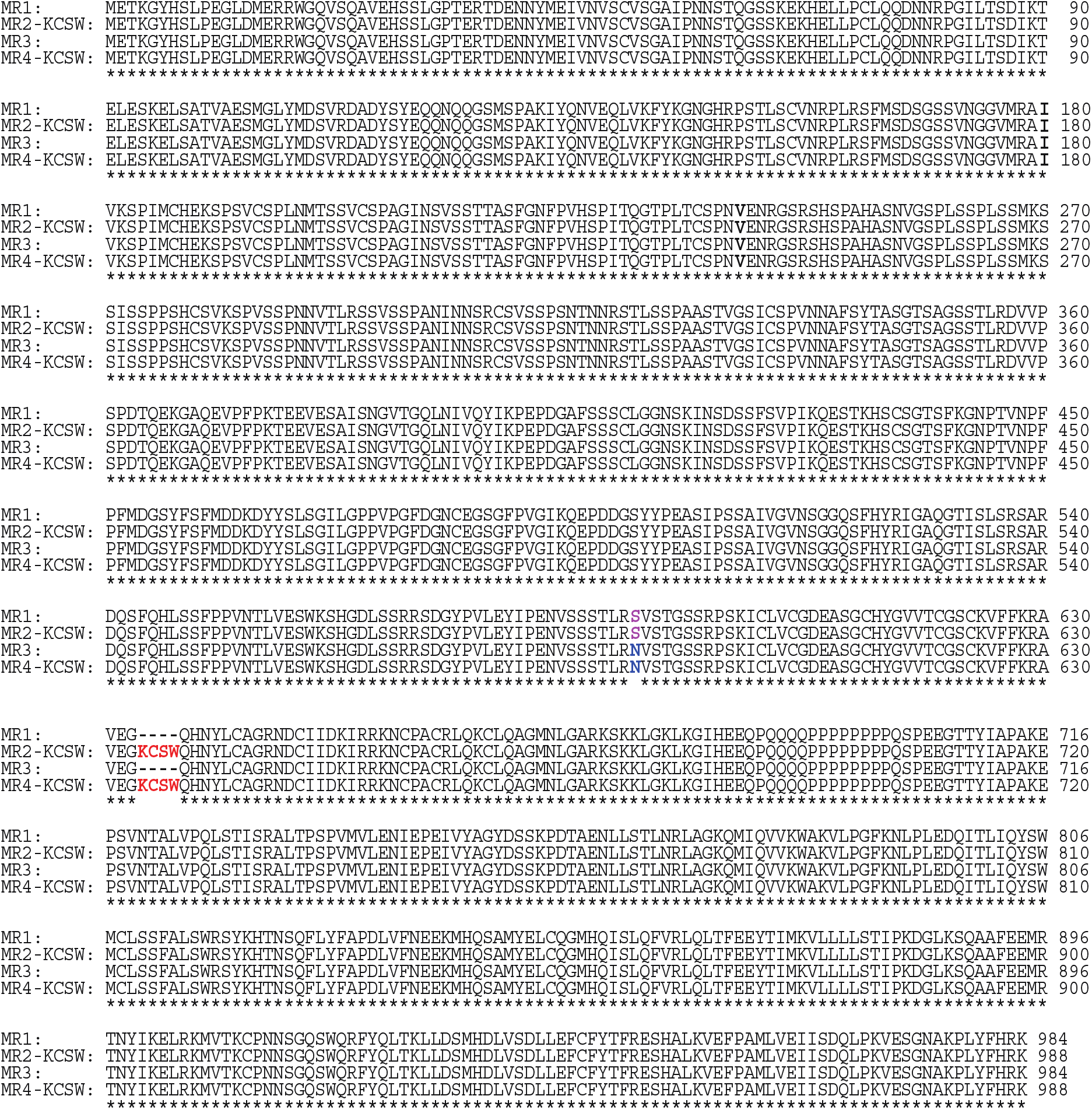
A multiple sequence alignment of four chimpanzee mineralocorticoid receptors. The four chimpanzee MRs were aligned with Clustal Omega [48]. In contrast to human MRs, chimpanzee MRs only contain an isoleucine at codon 180 and a valine at codon 241. Chimpanzees also contain an MR with either a serine or an asparagine at codon 591, while human MRs only contain a serine at codon 591.

### 3.3 Gorillas and Orangutans Have Two Full-Length MRs

Gorillas and orangutans each contain two full-length MRs: one MR contains 984 amino acids and one MR contains 988 amino acids (Figure 4). The MRs with 988 amino acids have the KCSW sequence in the DBD. All full-length gorilla MRs and full-length orangutan MRs have isoleucine-180 and valine-241 in the amino-terminal domain (NTD) (Figure 4). A gorilla MR or orangutan MR with either valine-180 and valine-241 or isoleucine-180 and alanine-241 in the NTD was not found in GenBank.

**Figure 4.**
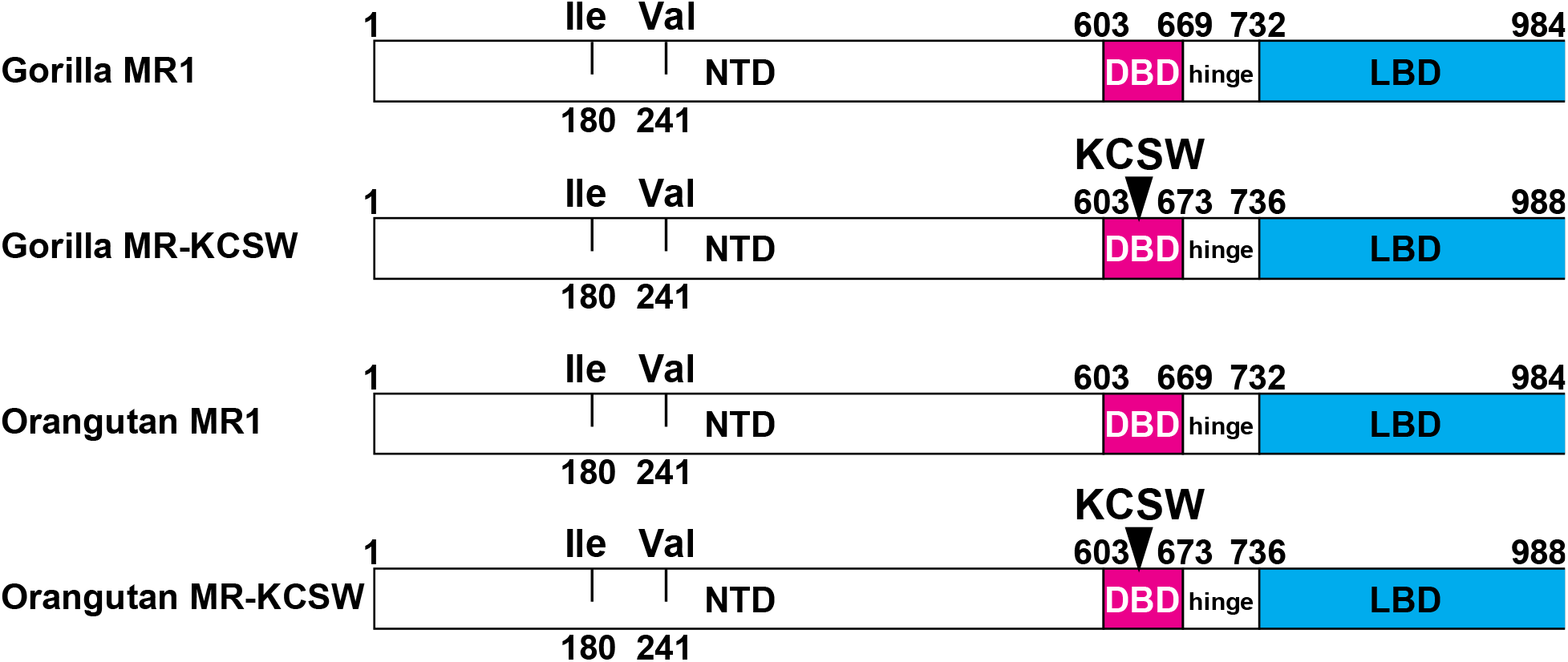
Comparison of the gorilla and orangutan MRs. The NTD of gorilla MRs and orangutan MRs conserve an isoleucine-180, valine-241 pair corresponding to isoleucine-180 and valine-241 in the NTD of human MR. A gorilla MR or an orangutan MR with either valine-180 and valine-241 or isoleucine-180 and alanine-241 in the NTD was not found in GenBank. Gorillas and orangutans also only contain an MR with a serine at codon 591, unlike chimpanzee MRs, which contain either serine at codon 591 or asparagine at codon 591. Human MRs only contain a serine at codon 591.

### 3.4. Evolutionary Divergence of Human and Chimpanzee Mineralocorticoid Receptors

Our analysis of human and chimpanzee MRs indicates that a human MR with either (Ile-180, Ala-241) or (Val-180, Val-241) in the NTD evolved after the divergence of humans and chimpanzees from a common ancestor. The physiological consequences of (Ile-180, Ala-241) or (Val-180, Val-241) in the NTD of human MR remain to be elucidated. Moreover, the absence of an asparagine at codon 591 in human MR is intriguing considering the presence of an asparagine at codon 591 in the MR in chimpanzees. As a first step, to determine if there are differences in transcriptional activation of these MRs by corticosteroids, we are cloning these human and chimpanzee MR genes and are screening them for transcriptional activation by aldosterone, cortisol, and other corticosteroids. We also will screen MR-GR dimers for activation by aldosterone and other corticosteroids [28,42–45,49], which is an important mechanism for regulating corticosteroid activity in the brain [15,17,44,50,51].

### 3.5. Evolution of the Mineralocorticoid Receptor DBD in Basal Terrestrial Vertebrates

The evolution of an MR with a DBD containing a four amino acid insert in human MR and other terrestrial MRs was surprising to us (Figure 1). We did not expect to find the KCSW insert in human MR at the position homologous to the position of TRQG in the DBD of lamprey CR because an insert at this position was not present in the DBD of elephant sharks and lungfish MRs (Figure 1). Moreover, the rest of the DBD sequence is highly conserved in terrestrial vertebrates (Figure 1) suggesting that the evolution of a four amino acid sequence in the DBD has an important function in terrestrial vertebrates. The evolution of the KCSR sequence into the DBD *Xenopus* MR (Figure 1) places the re-emergence of this motif in the MR close to the origin of terrestrial vertebrates, in which evolution of aldosterone synthesis in lungfish had an important role in the conquest of land by terrestrial vertebrates [4,6,52–58]. The evolution of KCSR in *Xenopus* MR in the homologous position to the DBD in human MR suggests that insertion of KCSR in *Xenopus* MR also may have been important early in the conquest of land by terrestrial vertebrates. Moreover, turtles contain an MR with KCSW in the homologous position in DBD (Figure 1) of human MR DBD, indicating that KCSW also has an ancient origin in MRs in terrestrial vertebrates. Indeed, the conservation of KCSW in the MR DBD in turtles and humans suggests an important function for KCSW in terrestrial vertebrates.

## Author Contributions

Author Contributions: Conceptualization, Yoshinao Katsu and Michael Baker; Data curation, Jiawen Zhang; Formal analysis, Yoshinao Katsu and Michael Baker; Investigation, Jiawen Zhang; Methodology, Yoshinao Katsu and Jiawen Zhang; Writing – original draft, Yoshinao Katsu and Michael Baker; Writing – review & editing, Yoshinao Katsu and Michael Baker.

## Institutional Review Board Statement

Not applicable.

## Data Availability Statement

The data presented in this study are available upon request from the corresponding author.

## References

1. Evans RM. The Steroid and Thyroid Hormone Receptor Superfamily. Science. 1988;240(4854):889–895. Doi:10.1126/Science.3283939.

2. Bridgham JT, Eick GN, Larroux C, et al. Protein Evolution by Molecular Tinkering: Diversification of the Nuclear Receptor Superfamily from a Ligand-Dependent Ancestor. PLoS Biol. 2010;8(10):E1000497. Published 2010 Oct 5. Doi:10.1371/Journal.Pbio.1000497.

3. Baker ME, Nelson DR, Studer RA. Origin of the Response to Adrenal and Sex Steroids: Roles of Promiscuity and Co-Evolution of Enzymes and Steroid Receptors. J Steroid Biochem Mol Biol. 2015;151:12–24. Doi:10.1016/j.Jsbmb.2014.10.020.

4. Baker ME. Steroid Receptors and Vertebrate Evolution. Mol Cell Endocrinol. 2019;496:110526. Doi:10.1016/j.Mce.2019.110526.

5. Evans RM, Mangelsdorf DJ. Nuclear Receptors, RXR, and the Big Bang. Cell. 2014 Mar 27;157(1):255–66. Doi: 10.1016/j.Cell.2014.03.012. PMID: 24679540; PMCID: PMC4029515.

6. Rossier BC, Baker ME, Studer RA. Epithelial Sodium Transport and Its Control by Aldosterone: The Story of Our Internal Environment Revisited. Physiol Rev. 2015;95(1):297–340. Doi:10.1152/Physrev.00011.2014.

7. Hanukoglu I, Hanukoglu A. Epithelial Sodium Channel (ENaC) Family: Phylogeny, Structure-Function, Tissue Distribution, and Associated Inherited Diseases. Gene. 2016;579(2):95–132. Doi:10.1016/j.Gene.2015.12.061.

8. Lifton RP, Gharavi AG, Geller DS. Molecular Mechanisms of Human Hypertension. Cell. 2001;104(4):545–556. Doi:10.1016/S0092-8674(01)00241-0.

9. Shibata S. 30 YEARS OF THE MINERALOCORTICOID RECEPTOR: Mineralocorticoid Receptor and NaCl Transport Mechanisms in the Renal Distal Nephron. J Endocrinol. 2017;234(1):T35–T47. Doi:10.1530/JOE-16-0669.

10. Grossmann C, Almeida-Prieto B, Nolze A, Alvarez de La Rosa D. Structural and Molecular Determinants of Mineralocorticoid Receptor Signalling. Br J Pharmacol. 2021 Nov 22. Doi: 10.1111/Bph.15746. Epub Ahead of Print. PMID: 34811739.

11. Hawkins UA, Gomez-Sanchez EP, Gomez-Sanchez CM, Gomez-Sanchez CE. The Ubiquitous Mineralocorticoid Receptor: Clinical Implications. Curr Hypertens Rep. 2012;14(6):573–580. Doi:10.1007/S11906-012-0297-0.

12. Jaisser F, Farman N. Emerging Roles of the Mineralocorticoid Receptor in Pathology: Toward New Paradigms in Clinical Pharmacology. Pharmacol Rev. 2016;68(1):49–75. Doi:10.1124/Pr.115.011106.

13. Funder J. 30 YEARS OF THE MINERALOCORTICOID RECEPTOR: Mineralocorticoid Receptor Activation and Specificity-Conferring Mechanisms: A Brief History. J Endocrinol. 2017 Jul;234(1):T17–T21. Doi: 10.1530/JOE-17-0119. Epub 2017 May 22. PMID: 28533421.

14. Gomez-Sanchez EP. Brain Mineralocorticoid Receptors in Cognition and Cardiovascular Homeostasis. Steroids. 2014 Dec;91:20–31. Doi: 10.1016/j.Steroids.2014.08.014. PMID: 25173821; PMCID: PMC4302001.

15. Joëls M, de Kloet ER. 30 YEARS OF THE MINERALOCORTICOID RECEPTOR: The Brain Mineralocorticoid Receptor: A Saga in Three Episodes. J Endocrinol. 2017 Jul;234(1):T49–T66. Doi: 10.1530/JOE-16-0660. PMID: 28634266.

16. Paul SN, Wingenfeld K, Otte C, Meijer OC. Brain Mineralocorticoid Receptor in Health and Disease: From Molecular Signalling to Cognitive and Emotional Function. Br J Pharmacol. 2022 Jul;179(13):3205–3219. Doi: 10.1111/Bph.15835. Epub 2022 Apr 7. PMID: 35297038; PMCID: PMC9323486.

17. Arriza JL, Simerly RB, Swanson LW, Evans RM. The Neuronal Mineralocorticoid Receptor as a Mediator of Glucocorticoid Response. Neuron. 1988 Nov;1(9):887–900. Doi: 10.1016/0896-6273(88)90136-5. PMID: 2856104.

18. Shimeld SM, Donoghue PC. Evolutionary Crossroads in Developmental Biology: Cyclostomes (Lamprey and Hagfish). Development. 2012 Jun;139(12):2091–9. Doi: 10.1242/Dev.074716. PMID: 22619386.

19. Close DA, Yun SS, McCormick SD, Wildbill AJ, Li W. 11-Deoxycortisol Is a Corticosteroid Hormone in the Lamprey. Proc Natl Acad Sci U S A. 2010 Aug 3;107(31):13942–7. Doi: 10.1073/Pnas.0914026107. Epub 2010 Jul 19. PMID: 20643930; PMCID: PMC2922276.

20. Thornton JW. Evolution of Vertebrate Steroid Receptors from an Ancestral Estrogen Receptor by Ligand Exploitation and Serial Genome Expansions. Proc Natl Acad Sci U S A 2001 May 8;98(10):5671–6. Doi: 10.1073/Pnas.091553298. Epub 2001 May 1. PMID: 11331759; PMCID: PMC33271.

21. Smith JJ, Kuraku S, Holt C, Sauka-Spengler T, Jiang N, Campbell MS, Yandell MD, Manousaki T, Meyer A, Bloom OE, Morgan JR, Buxbaum JD, Sachidanandam R, Sims C, Garruss AS, Cook M, Krumlauf R, Wiedemann LM, Sower SA, Decatur WA, Hall JA, Amemiya CT, Saha NR, Buckley KM, Rast JP, Das S, Hirano M, McCurley N, Guo P, Rohner N, Tabin CJ, Piccinelli P, Elgar G, Ruffier M, Aken BL, Searle SM, Muffato M, Pignatelli M, Herrero J, Jones M, Brown CT, Chung-Davidson YW, Nanlohy KG, Libants SV, Yeh CY, McCauley DW, Langeland JA, Pancer Z, Fritzsch B, de Jong PJ, Zhu B, Fulton LL, Theising B, Flicek P, Bronner ME, Warren WC, Clifton SW, Wilson RK, Li W. Sequencing of the Sea Lamprey (Petromyzon Marinus) Genome Provides Insights into Vertebrate Evolution. Nat Genet. 2013 Apr;45(4):415-21, 421e1-2. Doi: 10.1038/Ng.2568. Epub 2013 Feb 24. PMID: 23435085; PMCID: PMC3709584.

22. Baker ME, Katsu Y. 30 YEARS OF THE MINERALOCORTICOID RECEPTOR: Evolution of the Mineralocorticoid Receptor: Sequence, Structure and Function. J Endocrinol. 2017;234(1):T1–T16. Doi:10.1530/JOE-16-0661.

23. York JR, McCauley DW. The Origin and Evolution of Vertebrate Neural Crest Cells. Open Biol. 2020 Jan;10(1):190285. Doi: 10.1098/Rsob.190285. Epub 2020 Jan 29. PMID: 31992146; PMCID: PMC7014683.

24. Kuratani S. Evo-Devo Studies of Cyclostomes and the Origin and Evolution of Jawed Vertebrates. Curr Top Dev Biol. 2021;141:207–239. Doi: 10.1016/Bs.Ctdb.2020.11.011. Epub 2020 Dec 13. PMID: 33602489.

25. Janvier P. microRNAs Revive Old Views about Jawless Vertebrate Divergence and Evolution. Proc Natl Acad Sci U S A. 2010 Nov 9;107(45):19137–8. Doi: 10.1073/Pnas.1014583107. Epub 2010 Nov 1. PMID: 21041649; PMCID: PMC2984170.

26. Bridgham JT, Carroll SM, Thornton JW. Evolution of Hormone-Receptor Complexity by Molecular Exploitation. Science. 2006;312(5770):97–101. Doi:10.1126/Science.1123348.

27. Katsu Y, Lin X, Ji R, Chen Z, Kamisaka Y, Bamba K, Baker ME. N-Terminal Domain Influences Steroid Activation of the Atlantic Sea Lamprey Corticoid Receptor. J Steroid Biochem Mol Biol. 2023 Apr;228:106249. Doi: 10.1016/j.Jsbmb.2023.106249. Epub 2023 Jan 13. PMID: 36646152.

28. Katsu Y, Zhang J, Baker ME. Reduced Steroid Activation of Elephant Shark GR and MR after Inserting Four Amino Acids from the DNA-Binding Domain of Lamprey Corticoid Receptor-1. PLoS One. 2023 Aug 23;18(8):E0290159. Doi: 10.1371/Journal.Pone.0290159. PMID: 37611044; PMCID: PMC10446182.

29. Venkatesh B, Lee AP, Ravi V, Maurya AK, Lian MM, Swann JB, Ohta Y, Flajnik MF, Sutoh Y, Kasahara M, Hoon S, Gangu V, Roy SW, Irimia M, Korzh V, Kondrychyn I, Lim ZW, Tay BH, Tohari S, Kong KW, Ho S, Lorente-Galdos B, Quilez J, Marques-Bonet T, Raney BJ, Ingham PW, Tay A, Hillier LW, Minx P, Boehm T, Wilson RK, Brenner S, Warren WC. Elephant Shark Genome Provides Unique Insights into Gnathostome Evolution. Nature. 2014 Jan 9;505(7482):174–9. Doi: 10.1038/Nature12826. Erratum in: Nature. 2014 Sep 25;513(7519):574. Erratum in: Nature. 2020 Dec;588(7837):E15. PMID: 24402279; PMCID: PMC3964593.

30. Katsu Y, Kohno S, Oka K, et al. Transcriptional Activation of Elephant Shark Mineralocorticoid Receptor by Corticosteroids, Progesterone, and Spironolactone. Sci Signal. 2019;12(584):Eaar2668. Published 2019 Jun 4. Doi:10.1126/Scisignal.Aar2668, (n.d.).

31. Fonseca E, Machado AM, Vilas-Arrondo N, Gomes-Dos-Santos A, Veríssimo A, Esteves P, Almeida T, Themudo G, Ruivo R, Pérez M, Da Fonseca R, Santos MM, Froufe E, Román-Marcote E, Venkatesh B, Castro LFC. Cartilaginous Fishes Offer Unique Insights into the Evolution of the Nuclear Receptor Gene Repertoire in Gnathostomes. Gen Comp Endocrinol. 2020 Sep 1;295:113527. Doi: 10.1016/j.Ygcen.2020.113527. Epub 2020 Jun 8. PMID: 32526329.

32. Carroll SM, Bridgham JT, Thornton JW. Evolution of Hormone Signaling in Elasmobranchs by Exploitation of Promiscuous Receptors. Mol Biol Evol. 2008;25(12):2643–2652. Doi:10.1093/Molbev/Msn204.

33. Arriza JL, Weinberger C, Cerelli G, et al. Cloning of Human Mineralocorticoid Receptor Complementary DNA: Structural and Functional Kinship with the Glucocorticoid Receptor. Science. 1987;237(4812):268–275. Doi:10.1126/Science.3037703.

34. Fuller PJ. Novel Interactions of the Mineralocorticoid Receptor. Mol Cell Endocrinol. 2015 Jun 15;408:33–7. Doi: 10.1016/j.Mce.2015.01.027. Epub 2015 Feb 7. PMID: 25662276.

35. Hellal-Levy C, Couette B, Fagart J, Souque A, Gomez-Sanchez C, Rafestin-Oblin M. Specific Hydroxylations Determine Selective Corticosteroid Recognition by Human Glucocorticoid and Mineralocorticoid Receptors. FEBS Lett. 1999;464(1-2):9–13. Doi:10.1016/S0014-5793(99)01667-1.

36. Geller DS, Farhi A, Pinkerton N, et al. Activating Mineralocorticoid Receptor Mutation in Hypertension Exacerbated by Pregnancy. Science. 2000;289(5476):119–123. Doi:10.1126/Science.289.5476.119.

37. Altschul SF, Gish W, Miller W, Myers EW, Lipman DJ. Basic Local Alignment Search Tool. J Mol Biol. 1990 Oct 5;215(3):403–10. Doi: 10.1016/S0022-2836(05)80360-2. PMID: 2231712.

38. Bloem LJ, Guo C, Pratt JH. Identification of a Splice Variant of the Rat and Human Mineralocorticoid Receptor Genes. J Steroid Biochem Mol Biol. 1995 Nov;55(2):159–62. Doi: 10.1016/0960-0760(95)00162-s. PMID: 7495694.

39. Wickert L, Watzka M, Bolkenius U, Bidlingmaier F, Ludwig M. Mineralocorticoid Receptor Splice Variants in Different Human Tissues. Eur J Endocrinol. 1998 Jun;138(6):702–4. Doi: 10.1530/Eje.0.1380702. PMID: 9678540.

40. Wickert L, Selbig J, Watzka M, Stoffel-Wagner B, Schramm J, Bidlingmaier F, Ludwig M. Differential mRNA Expression of the Two Mineralocorticoid Receptor Splice Variants within the Human Brain: Structure Analysis of Their Different DNA Binding Domains. J Neuroendocrinol. 2000 Sep;12(9):867–73. Doi: 10.1046/j.1365-2826.2000.00535.x. PMID: 10971811.

41. De Kloet ER, Joëls M. Brain Mineralocorticoid Receptor Function in Control of Salt Balance and Stress-Adaptation. Physiol Behav. 2017 Sep 1;178:13–20. Doi: 10.1016/j.Physbeh.2016.12.045. Epub 2017 Jan 13. PMID: 28089704.

42. Liu W, Wang J, Sauter NK, Pearce D. Steroid Receptor Heterodimerization Demonstrated in Vitro and in Vivo. Proc Natl Acad Sci U S A. 1995 Dec 19;92(26):12480–4. Doi: 10.1073/Pnas.92.26.12480. PMID: 8618925; PMCID: PMC40381.

43. Trapp T, Holsboer F. Heterodimerization between Mineralocorticoid and Glucocorticoid Receptors Increases the Functional Diversity of Corticosteroid Action. Trends Pharmacol Sci. 1996 Apr;17(4):145–9. Doi: 10.1016/0165-6147(96)81590-2. PMID: 8984741.

44. Mifsud KR, Reul JM. Acute Stress Enhances Heterodimerization and Binding of Corticosteroid Receptors at Glucocorticoid Target Genes in the Hippocampus. Proc Natl Acad Sci U S A. 2016 Oct 4;113(40):11336–11341. Doi: 10.1073/Pnas.1605246113. Epub 2016 Sep 21. PMID: 27655894; PMCID: PMC5056104.

45. Pooley JR, Rivers CA, Kilcooley MT, Paul SN, Cavga AD, Kershaw YM, Muratcioglu S, Gursoy A, Keskin O, Lightman SL. Beyond the Heterodimer Model for Mineralocorticoid and Glucocorticoid Receptor Interactions in Nuclei and at DNA. PLoS One. 2020 Jan 10;15(1):E0227520. Doi: 10.1371/Journal.Pone.0227520. PMID: 31923266; PMCID: PMC6953809.

46. Kiilerich P, Triqueneaux G, Christensen NM, et al. Interaction between the Trout Mineralocorticoid and Glucocorticoid Receptors in Vitro. J Mol Endocrinol. 2015;55(1):55- Doi:10.1530/JME-15-0002.

47. Altschul SF, Madden TL, Schäffer AA, Zhang J, Zhang Z, Miller W, Lipman DJ. Gapped BLAST and PSI-BLAST: A New Generation of Protein Database Search Programs. Nucleic Acids Res. 1997 Sep 1;25(17):3389–402. Doi: 10.1093/Nar/25.17.3389. PMID: 9254694; PMCID: PMC146917.

48. Sievers F, Wilm A, Dineen D, Gibson TJ, Karplus K, Li W, Lopez R, McWilliam H, Remmert M, Söding J, Thompson JD, Higgins DG. Fast, Scalable Generation of High-Quality Protein Multiple Sequence Alignments Using Clustal Omega. Mol Syst Biol. 2011 Oct 11;7:539. Doi: 10.1038/Msb.2011.75. PMID: 21988835; PMCID: PMC3261699.

49. Clarisse D, Prekovic S, Vlummens P, Staessens E, Van Wesemael K, Thommis J, Fijalkowska D, Acke G, Zwart W, Beck IM, Offner F, De Bosscher K. Crosstalk between Glucocorticoid and Mineralocorticoid Receptors Boosts Glucocorticoid-Induced Killing of Multiple Myeloma Cells. Cell Mol Life Sci. 2023 Aug 14;80(9):249. Doi: 10.1007/S00018-023-04900-x. PMID: 37578563; PMCID: PMC10425521.

50. De Kloet ER. Brain Mineralocorticoid and Glucocorticoid Receptor Balance in Neuroendocrine Regulation and Stress-Related Psychiatric Etiopathologies. Curr Opin Endocr Metab Res. 2022 Jun;24:100352. Doi: 10.1016/j.Coemr.2022.100352. PMID: 38037568; PMCID: PMC10687720.

51. De Kloet ER, de Kloet SF, de Kloet CS, de Kloet AD. Top-down and Bottom-up Control of Stress-Coping. J Neuroendocrinol. 2019 Mar;31(3):E12675. Doi: 10.1111/Jne.12675. Epub 2019 Feb 1. PMID: 30578574; PMCID: PMC6519262.

52. Katsu Y, Oana S, Lin X, Hyodo S, Baker ME. Aldosterone and Dexamethasone Activate African Lungfish Mineralocorticoid Receptor: Increased Activation after Removal of the Amino-Terminal Domain. J Steroid Biochem Mol Biol. 2022 Jan;215:106024. Doi: 10.1016/j.Jsbmb.2021.106024. Epub 2021 Nov 10. PMID: 34774724.

53. Uchiyama M, Maejima S, Yoshie S, Kubo Y, Konno N, Joss JM. The Epithelial Sodium Channel in the Australian Lungfish, Neoceratodus Forsteri (Osteichthyes: Dipnoi). Proc Biol Sci. 2012;279(1748):4795–4802. Doi:10.1098/Rspb.2012.1945.

54. Joss JM. Lungfish Evolution and Development. Gen Comp Endocrinol. 2006 Sep 15;148(3):285–9. Doi: 10.1016/j.Ygcen.2005.10.010. Epub 2005 Dec 7. PMID: 16337631.

55. Biscotti MA, Gerdol M, Canapa A, Forconi M, Olmo E, Pallavicini A, Barucca M, Schartl M. The Lungfish Transcriptome: A Glimpse into Molecular Evolution Events at the Transition from Water to Land. Sci Rep. 2016 Feb 24;6:21571. Doi: 10.1038/Srep21571. PMID: 26908371; PMCID: PMC4764851.

56. McCormick SD, Bradshaw D. Hormonal Control of Salt and Water Balance in Vertebrates. Gen Comp Endocrinol. 2006 May 15;147(1):3–8. Doi: 10.1016/j.Ygcen.2005.12.009. Epub 2006 Feb 2. PMID: 16457828.

57. Fishman AP. Homer W. SMITH (1895-1962). Circulation. 1962 Nov;26:984–5. Doi: 10.1161/01.Cir.26.5.984. PMID: 13945307.

58. Rossier BC. Osmoregulation during Long-Term Fasting in Lungfish and Elephant Seal: Old and New Lessons for the Nephrologist. Nephron. 2016;134(1):5–9. Doi: 10.1159/000444307. Epub 2016 Feb 23. PMID: 26901864.

